# Online Closed-Loop Real-Time tES-fMRI for Brain Modulation: Feasibility, Noise/Safety and Pilot Study

**DOI:** 10.1101/2021.04.10.439268

**Authors:** Beni Mulyana, Aki Tsuchiyagaito, Jared Smith, Masaya Misaki, Rayus Kuplicki, Ghazaleh Soleimani, Ashkan Rashedi, Duke Shereen, Til Ole Bergman, Samuel Cheng, Martin Paulus, Jerzy Bodurka, Hamed Ekhtiari

## Abstract

Recent studies suggest that transcranial electrical stimulation (tES) can be performed during functional magnetic resonance imaging (fMRI). The novel approach of using concurrent tES-fMRI to modulate and measure targeted brain activity/connectivity may provide unique insights into the causal interactions between the brain neural responses and psychiatric/neurologic signs and symptoms, and importantly, guide the development of new treatments. However, tES stimulation parameters to optimally influence the underlying brain activity in health and disorder may vary with respect to phase, frequency, intensity and electrode’s montage. Here, we delineate how a closed-loop tES-fMRI study of frontoparietal network modulation can be designed and performed. We also discuss the challenges of running a concurrent tES-fMRI, describing how we can distinguish clinically meaningful physiological changes caused by tES from tES-related artifacts. There is a large methodological parameter space including electrode types, electrolytes, electrode montages, concurrent tES-fMRI hardware, online fMRI processing pipelines and closed-loop optimization algorithms that should be carefully selected for closed-loop tES-fMRI brain modulation. We also provide technical details on how safety and quality of tES-fMRI settings can be tested, and how these settings can be monitored during the study to ensure they do not exceed safety standards. The initial results of feasibility and applicability of closed-loop tES-fMRI are reported and potential hypotheses for the outcomes are discussed.

**Highlight points:**

- Methodological details of a closed-loop tES-fMRI study protocol are provided.
- The protocol is performed successfully on a frontoparietal network without side-effects.
- The temperature of electrodes in concurrent tES-fMRI remains in the safe range.
- Properly setup concurrent tES does not introduce MRI artifacts and noise.
- Simplex optimizer could be used to find an optimal tES stimulation parameter.

## Introduction

Transcranial electrical stimulation (tES) provides electric current stimulation over the scalp to modulate specific brain regions’ neural activity or their functional connectivity (Bikson et al., 2019). This method can be concurrently combined with functional magnetic resonance imaging (fMRI). Such tES-fMRI combination has several technical advantages (Saiote, Turi, Paulus, & Antal, 2013; Williams et al., 2017) compared with: 1) sequential fMRI-tES-fMRI in terms of the ability to investigate ongoing brain activity, and 2) simultaneous tES-electroencephalography (EEG) in terms of higher spatial resolution and fewer problems with stimulation artifacts. A major advantage of concurrent tES with fMRI is that we can stimulate several regions of the brain by tES (i.e. two nodes of a network with conventional or high definition (HD) electrode montages) and evaluate its online stimulation effect by fMRI to reveal associations between brain stimulation and whole-brain activity/connectivity (Bächinger et al., 2017; Cabral-Calderin, Williams, Opitz, Dechent, & Wilke, 2016; Violante et al., 2017; Vosskuhl, Huster, & Herrmann, 2016).

tES is a non-invasive brain stimulation (NIBS) technique including direct (tDCS), alternating current (tACS) and random noise stimulation (tRNS) (Bikson et al., 2019). Although all tES methods can target large scale brain networks, tACS has the unique potential to modulate oscillations within or between the large scale brain networks using alternating currents at a chosen frequency and phase difference between network to interact with synchronization-based functional connectivity (Ruffini, Fox, Ripolles, Miranda, & Pascual-Leone, 2014). The blood oxygenation level dependent (BOLD) fMRI signal relies on the blood flow response to brain neuronal activity, which is much slower than the electrophysiological activity of the individual neuron. The BOLD response starts to increase a few seconds after the respective change in neural activation. However, the presence of correlated BOLD signal fluctuations at rest (e.g., resting state networks) across brain areas may result from oscillatory synchronization facilitating communication between those regions (Buzsáki & Draguhn, 2004; Canolty & Knight, 2010), as supported by findings from concurrent EEG-fMRI studies demonstrating the association of electrophysiological neuronal oscillation as measured by EEG and BOLD signal large spatial scale synchronization (Mantini, Perrucci, Del Gratta, Romani, & Corbetta, 2007; Whitman, Ward, & Woodward, 2013; Yuan et al., 2016; Yuan, Zotev, Phillips, Drevets, & Bodurka, 2012). Therefore, fMRI functional connectivity across various brain regions may serve as a proxy-marker to measure internal co-oscillatory electrophysiological synchronization of those regions. External oscillatory stimulation (e.g., at frequency range matching EEG rhythmic activity: 0.1-40Hz or even higher) above several cortical regions using multi-site tACS has been demonstrated to increase internal oscillatory synchronization and functional connectivity between brain regions as well as cognitive function (Cabral-Calderin et al., 2016; Kuo & Nitsche, 2012; Moisa, Polania, Grueschow, & Ruff, 2016; Violante et al., 2017; Weinrich et al., 2017a; Williams et al., 2017; Zoefel, Archer-Boyd, & Davis, 2018). However, determining the ideal configuration of a multi-site tACS system aimed at modulating brain networks is complex as the effects of tACS are highly dependent on the stimulation parameters such as electric current stimulation intensity, frequency, and inter-regional phase differences, selection of electrode locations and individual differences in brain structure (Antal & Paulus, 2013). For example, a plausible range of stimulation frequencies (0.1-100 Hz) and phases differences (0-359°) between stimulation sites (Lorenz et al., 2019) result in a wide-range of possibilities. Establishing optimization algorithms would aid the clinical application of tACS. More precisely, an online fMRI measurement will enable us to establish empirically an optimization algorithm by identifying the stimulation parameters (i.e., frequency and phase differences in this study) which maximize the targeted brain network activity/connectivity (i.e., temporal correlations between BOLD signal changes in two target regions). Moreover, applying multi-electrode configurations, or high definition montages (Datta et al., 2009) has been shown to result in more focal electric field distribution patterns (Alam, Truong, Khadka, & Bikson, 2016; Villamar et al., 2013), which will also allow unique combinations of electrode locations combined with optimized stimulation parameters to more focally target specific cortical regions (Dmochowski, Datta, Bikson, Su, & Parra, 2011).

We report our recently developed MRI-conditional high-definition tACS (HD-tACS) setup using two sites 4□1 ring montages for frontoparietal synchronization (FPS) (Saturnino, Madsen, Siebner, & Thielscher, 2017) in combination with an optimization algorithm to achieve, for the first time, fully closed-loop tACS-fMRI. We provide the details of the online FPS closed-loop tACS-fMRI experimental protocol to test the efficacy of this intervention and sample data, expected outcomes and hypotheses. Moreover, we show how the effect of tACS on brain activity, as inferred from the BOLD signal, can be measured and validated, while excluding technical artifacts in fMRI signal related to tACS. We discuss safety aspects (i.e., temperature under electrodes and patient comfort, sensation and side effects) of closed-loop tACS-fMRI settings.

## Online frontoparietal synchronization closed-loop tACS-fMRI protocol study design

Frontoparietal connectivity within the executive control network (ECN) is considered one of the main therapeutic targets for network-based brain stimulation in different psychiatric disorders, e.g., depression and substance use disorders (Ekhtiari, Nasseri, Yavari, Mokri, & Monterosso, 2016; Fischer, Keller, & Etkin, 2016). Here, we propose an online frontoparietal stimulation (FPS) closed-loop tACS-fMRI protocol, which can be utilized for future clinical and experimental studies. The purpose of this protocol is to examine the effectiveness of online FPS with tACS-fMRI (FPS session) and to determine the optimal frequency and phase difference of multi-site tACS to enhance frontoparietal connectivity during stimulation. Participants are asked to answer several self-surveys before and after the FPS session to evaluate the feasibility and the side-effect of FPS session (Figure 1A). Participants then undergo a 2-back task as a cognitive function baseline (Jaeggi, Buschkuehl, Perrig, & Meier, 2010; Owen, McMillan, Laird, & Bullmore, 2005) before the FPS session. After the baseline measurement of the cognitive function, they undergo an anatomical scan, a first resting scan, a first FPS with a 2-back task (training 1), a second FPS with a 2-back task (training 2), and a second resting scan. During the two training FPS sessions, participants are randomly assigned to either an optimized (experimental) group or a control group. For the optimized parameters (experimental group), the optimizer searches the tACS parameter that gives the highest target frontoparietal connectivity, otherwise, control parameters (control group) search for the worst or lowest target frontoparietal connectivity during the training runs. Then the participant undergoes a testing FPS scan, in which they are stimulated with the optimized (experimental group) or control parameters (control group) with a 2-back task (testing). Finally, participants undergo a third resting scan (Figure 1B).

**Figure 1.**
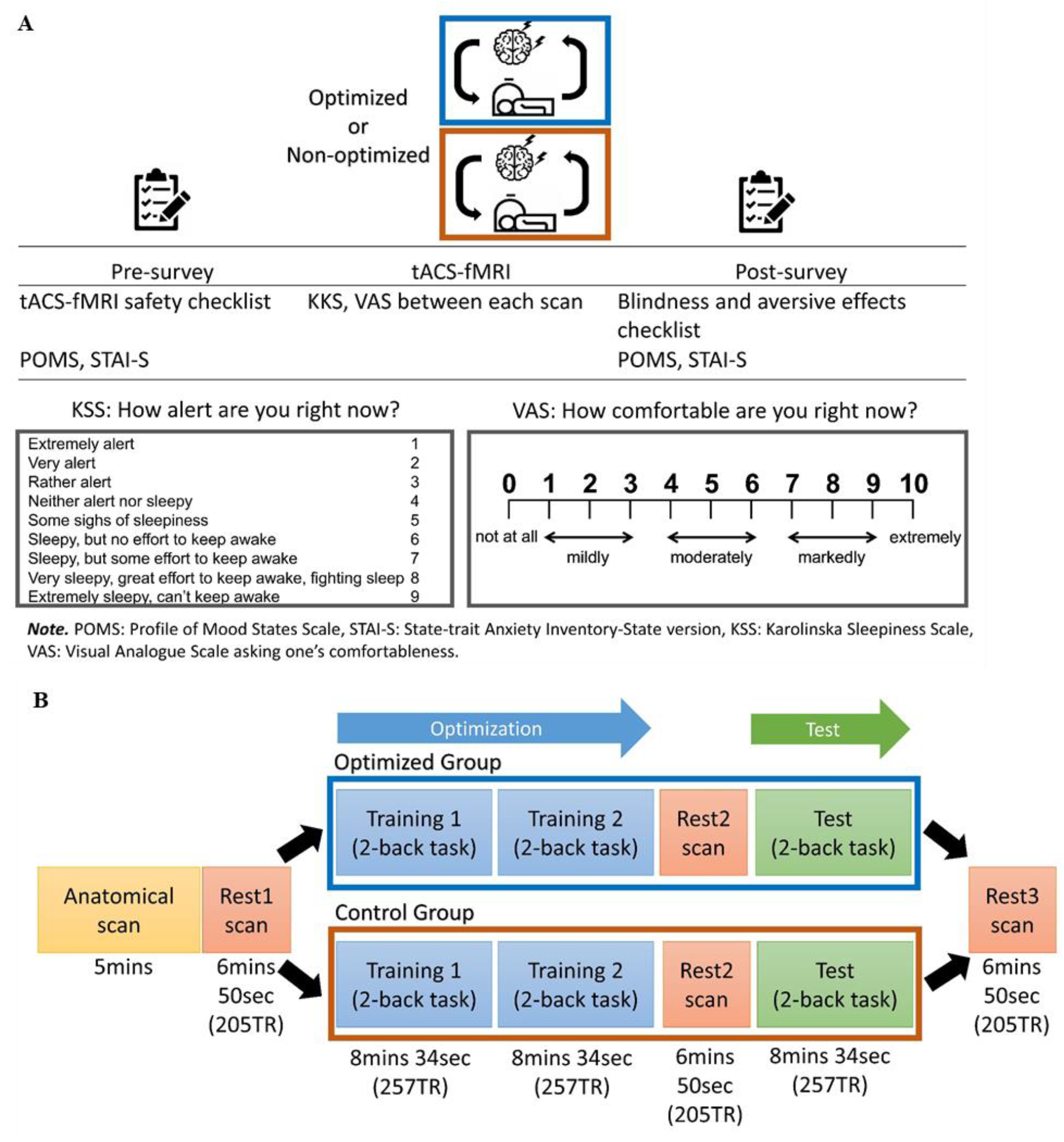
Study design. **A**) an entire session of the online frontoparietal synchronization closed-loop tACS-fMRI protocol; **B)** a detailed overview of the closed-loop tACS-fMRI part. TR = time repetition.

From 2-back task results as cognitive performance measurement, we derive accuracy and response time of correct answers pre- and post-FPS session. Details of each training scan and testing scan are described below.

The tACS stimulation was applied using an MRI-compatible Starstim AC-Stimulator (https://www.neuroelectrics.com/products/starstim/starstim-r32/). FPS session targeted the right middle frontal gyrus (R-DLPFC,) and right inferior parietal cortex (R-IPC) as the important nodes of the frontoparietal network, approximated by electrode positions F4 and P4 of the 10-20 EEG system. The electrode is a modification of MRI Sponstim (model: NE026MRI, brand: Neuroelectrics), placed inside the next-generation (NG) Pistim’s shell (model:NE029, brand: Neuroelectrics) with the metal part (Ag/AgCl) of the shell removed. This modification creates an MRI-compatible electrode (circular pad radius = 1cm with carbon rubber as electrode pad) and combined with conductive gel/paste (model: Abralyt HiCl, brand: Easycap), it improves contact conductivity between the scalp and the carbon rubber pad. We use textile caps with holes indicating places for electrode positioning (model: Neoprene Headcap/ NE019, brand: Neuroelectrics).

The stimulation is divided into two stages (training and testing). The training stage determines the optimum frequency and phase difference parameters that produce the highest frontoparietal network connectivity while subjects perform a cognitive task (here, a 2-back task). The training run is divided into 15 blocks (Figure 2A) where each block consists of 20 seconds tACS with parameters (frequency and phase) derived from the Simplex optimizer rules. This is followed by 10 seconds of rest. During each block, the Simplex optimizer searches the optimized parameter of the combination of frequency and phase from the parameters’ field (i.e., two-dimensional parameters’ field of the frequency (1-150Hz) and phase difference (0–359°)) based on the fMRI frontoparietal functional connectivity measurements.

**Figure 2.**
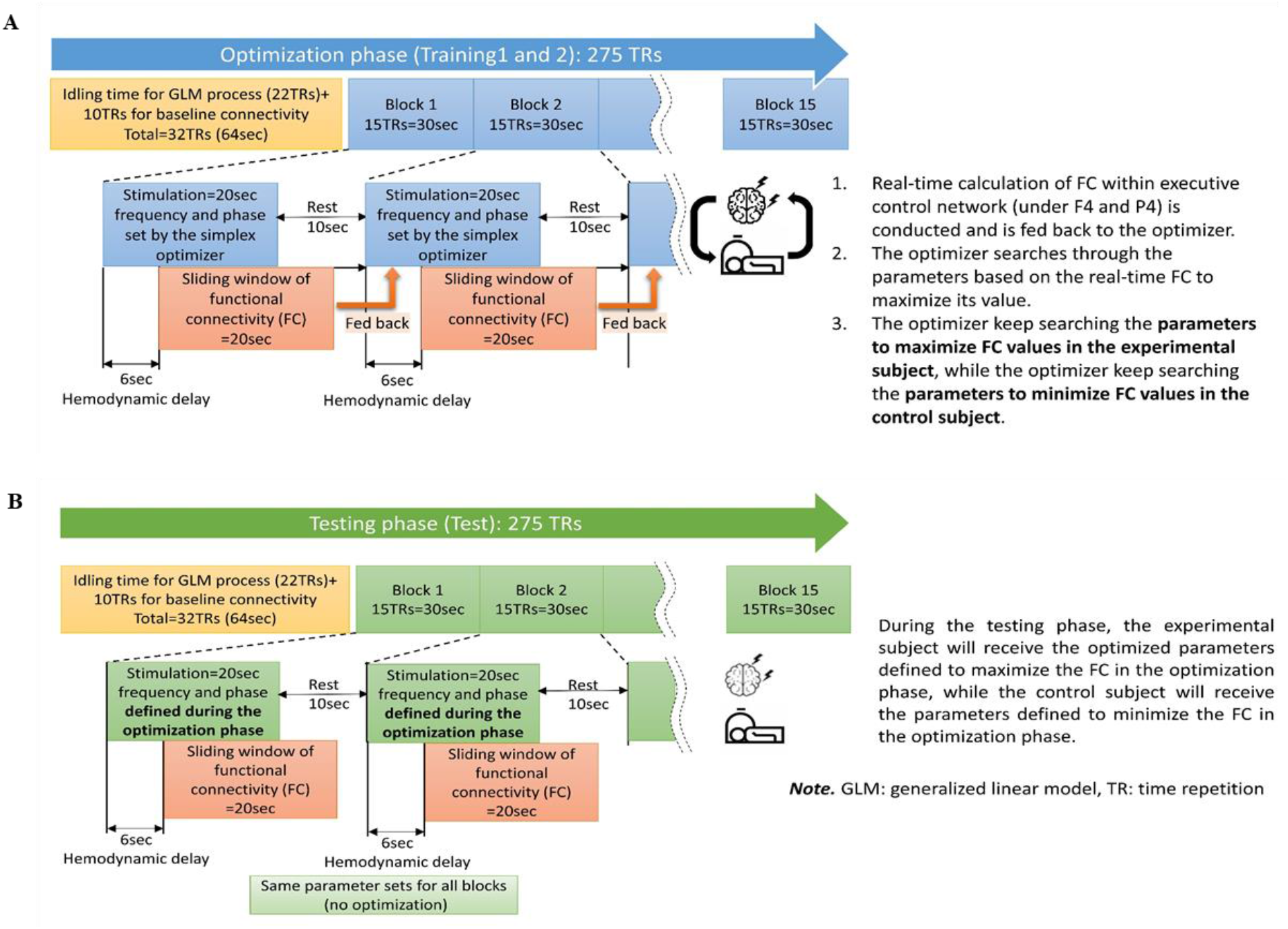
Overview of the optimization approach. **A)** Training 1 and 2 runs in order to find optimal tACS parameters; **B)** Testing run to test the optimal parameters which is found by training run.

The optimizer uses the Simplex algorithm and the Nelder-Mead technique. Details of this method are explained in (Mathews & Fink, 2004; Nelder & Mead, 1965; Singer & Nelder, 2009). Briefly, it is a simple optimization algorithm seeking the vector of parameters, which, in this study, are frequency and phase difference between F4 and P4 sites, that correspond to the maximum fMRI frontoparietal functional connectivity (more details are available in section 2.5). The sliding-window fMRI functional connectivity response is calculated in real-time and analyzed by the Simplex optimizer to predict what frequency and phase cause the highest increase in frontoparietal functional connectivity.

The MRI anatomical image was aligned to one fMRI echo planar image (EPI) image on Rest1, and then the frontoparietal reference mask in MNI space to that result to create an individual mask to calculate frontoparietal connectivity. Before the first stimulation block, there is an idling time for a general linear model (GLM) process to regress noise from the fMRI signal for 22 TRs (1 TR = 2 seconds, 22 TRs = 44 seconds) and 10 TR (20 seconds) for baseline functional connectivity calculations. The optimization phase takes approximately 9 minutes and is repeated twice (training 1 and 2), to reduce the burden for the participant and give them a short break between scans. The testing phase (testing run) tests the optimized parameters’ ability to modulate the ECN and compares them to the results when using control parameters. The testing run is similar to the training, which is divided into 15 blocks (Figure 2B).

This testing run lasts approximately 9 mins as the training runs. However, during the testing run, we do not use the optimizer to calculate optimized parameters but only use the optimized parameters obtained from training 1 and 2 or the respective control parameters. Participants perform a 2-back task during the training 1, 2, and testing stages (details are explained in 2.6).

### 2.1 Online closed-loop tACS-fMRI equipment settings

The battery-driven tACS device is positioned outside the magnetic field in the operator room (Figure 3).

**Figure 3.**
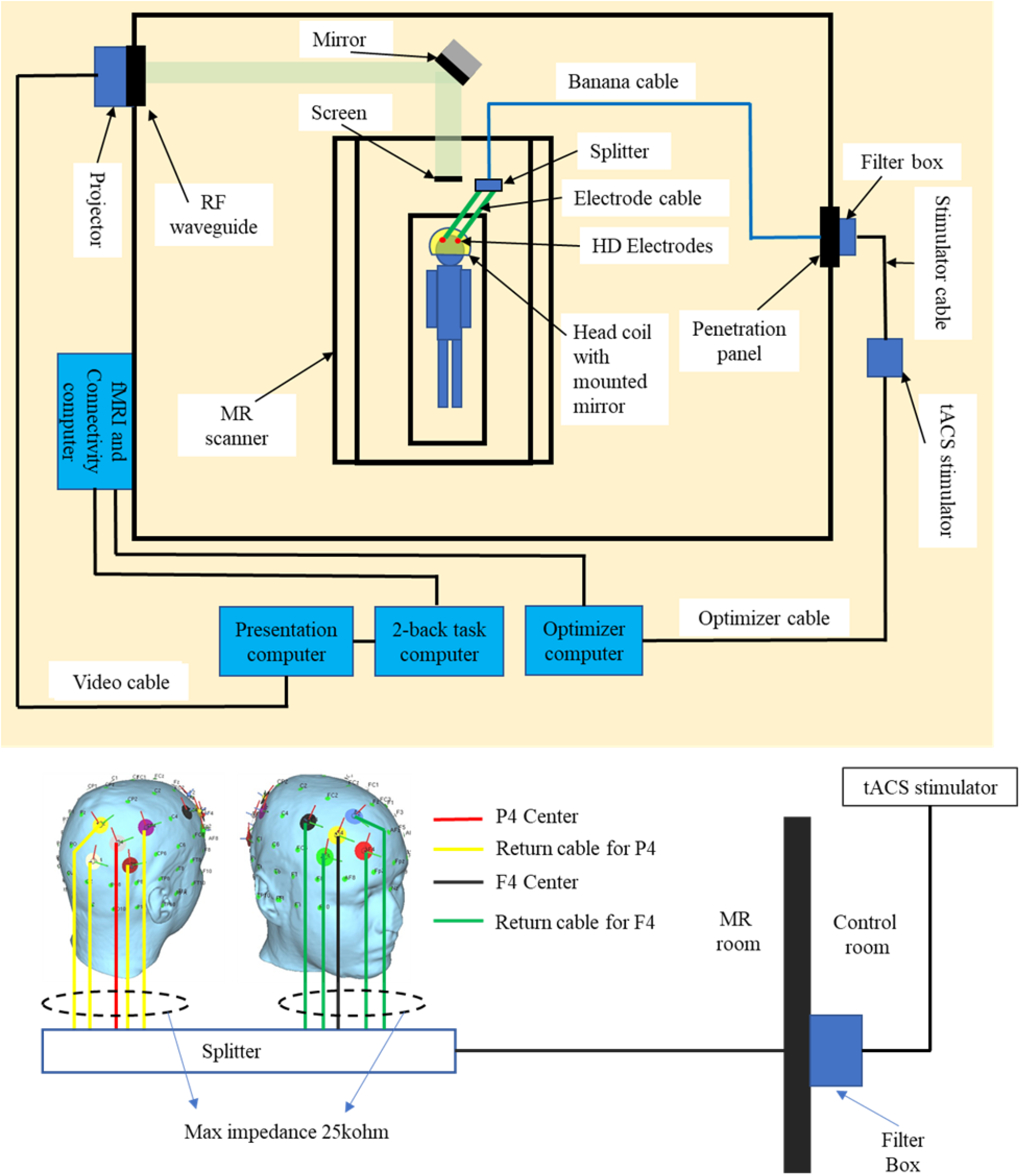
Closed-loop tES-fMRI setup. The participant is capped with frontoparietal 10HD electrodes, then laying down inside MRI room to get tACS stimulation concurrent with fMRI scanning. During fMRI scanning, fMRI connectivity computer sends frontoparietal connectivity to 2-back task computer and Optimizer computer. 2-back task computer connected to the presentation computer to display 2-back task on the screen inside MRI room for the participant. The optimizer calculates the optimal tACS parameters for improving participant frontoparietal functional connectivity. Then, the optimizer sends the tACS parameters through the optimizer cable to the tACS device. The tACS device is connected to the filter box that attached on the penetration panel using a stimulator cable. Then, the filter box is connected through a banana cable to the participant frontoparietal sites via 10HD electrodes to give a stimulation. TR = time repetition.

Stimulation is channeled into the scanner bore via a filter box (MECMRI-Series, 2018) attached to the penetration panel that filters out radio frequency (RF) noise (7–1000 MHz) and high magnetic fields from the scanner. The stimulation electrodes, which are built to be safe in an MRI environment, are positioned at F4 and P4 in the 10-20 system, and four electrodes surrounding F4 and P4 as a return for their center (F4 or P4). Return-electrodes placement for F4 and P4 sites are designed to be at an equal center-return distance (3cm) in order to avoid gel bridging (short circuit) and to avoid the electrical shunt effect in anti-phase condition. The electrical shunt effect is explained in section 2.3. Return-electrode coordinates for the F4 site are: R_F1_ = [37.99, 75.64, 22.32], R_F2_ = [67.91, 48.88, 25.73], R_F3_ = [52.23, 35.78, 60.45], and R_F4_ = [22.31, 62.54, 57.04] (Figure 4A) in the position coordinates of the standard BioSemi headcaps with circumference = 550mm (https://www.biosemi.com/headcap.htm). Return-electrode coordinates for P4 site are:(RP1 = [22.12, −60.72, 59.04], R_P2_ = [50.21, −35.60, 62.24], R_P3_ = [69.58, −43.35, 30.70], and R_P4_ = [48.22, −71.16, 16.55] (Figure 4B) in the position coordinates of the standard BioSemi headcaps with circumference = 550mm). Care is taken to avoid placing the return electrodes on PO4 l, because this electrode site would result in uncomfortable pressure on the back of the subject’s head when laying on the MRI table (the red area in Figure 4A).

**Figure 4.**
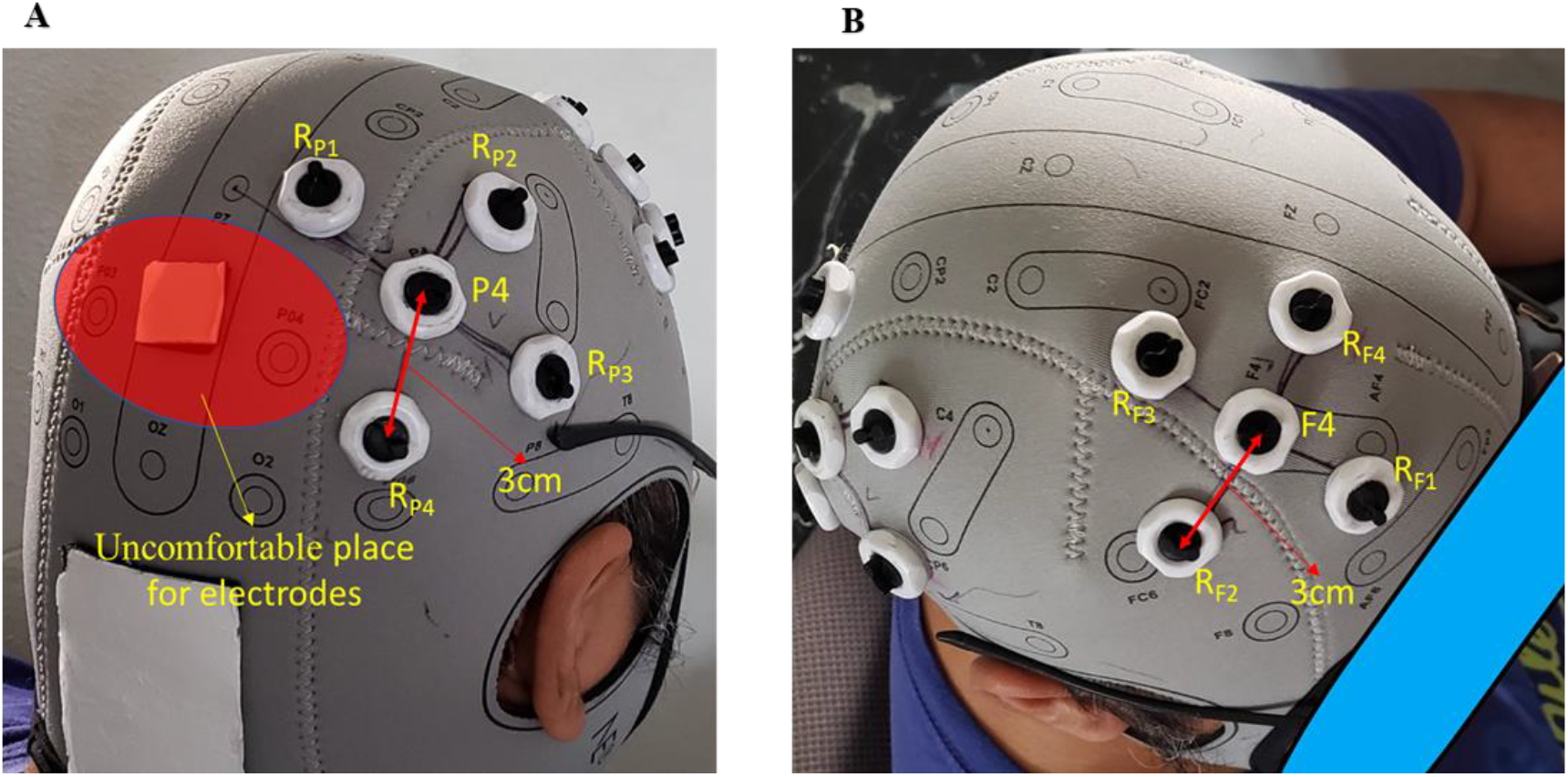
Overview of 10 HD electrodes for this study. **A)** Montage of 10 HD electrodes on the parietal site with equidistant (3 cm) center and return electrodes. The red highlighted area is a subject uncomfortable area where we need to avoid placing the electrodes due to the constant head pressure when subjects lie down on the MRI table; **B)** Montage of 10 HD electrodes on the frontal site with equidistant (3 cm) center and return electrodes.

The coordinates of the highest electric field on frontal and parietal montage sites using SimNIBS software for E-field simulation (Saturnino, Madsen, & Thielscher, 2019) are assigned as the center of the frontoparietal reference mask in the frontal site (MNI coordinates = [−45, 49, 27] and radius = 10mm) and in the parietal site (MNI coordinates = [−45, −75, 46] and radius = 10mm). fMRI connectivity provides the feedback for optimizing the tACS parameters (frequency and phase difference between F4 and P4) to increase F4-P4 connectivity. After convergence or a maximum of 30 iterations on training 1 and 2, the optimized protocol to induce maximum online frontoparietal connectivity will be determined (Kingma & Ba, 2015; Lancaster, Lorenz, Leech, & Cole, 2018; Lorenz, Hampshire, & Leech, 2017; Lorenz et al., 2016; Monti et al., 2017; Nelli & Nelli, 2018; Shukla, 2018).

### 2.2 Electrodes

Currently, the Starstim R32 tACS device uses the rubber electrode embedded in a sponge pocket with saline solution as a conductive material between the electrode and scalp. Although this electrode solution is more comfortable for participants compared to conductive gel, using saline solution has many disadvantages, such as; (1) saline solution evaporates quickly, and it would be difficult to maintain safe and low impedance during the long duration of experiments; (2) saline solution is easy to spread out and has a greater risk to make short circuits between electrodes, which will not ensure the accurate stimulation over the desired sites of cortical area; (3) the sponge is made of textile sponge and the contact with the carbon rubber could be loose. To overcome those potential disadvantages of saline solution and also to take advantages of the focality of HD electrodes, we created MRI compatible rubber HD electrodes (circular pad with radius 1cm and 1mm thickness, electrode material: carbon rubber and plastic shell) (Figure 5A) and use Abralyt HiCl highly conductive gel (Piervirgili, Petracca, & Merletti, 2014) as a conductor. Our electrode shell construction has a dome structure so that it avoids gel spreading out over the scalp and will be a better setting compared with electrodes embedded in sponge pockets soaked with saline solution.

**Figure 5.**
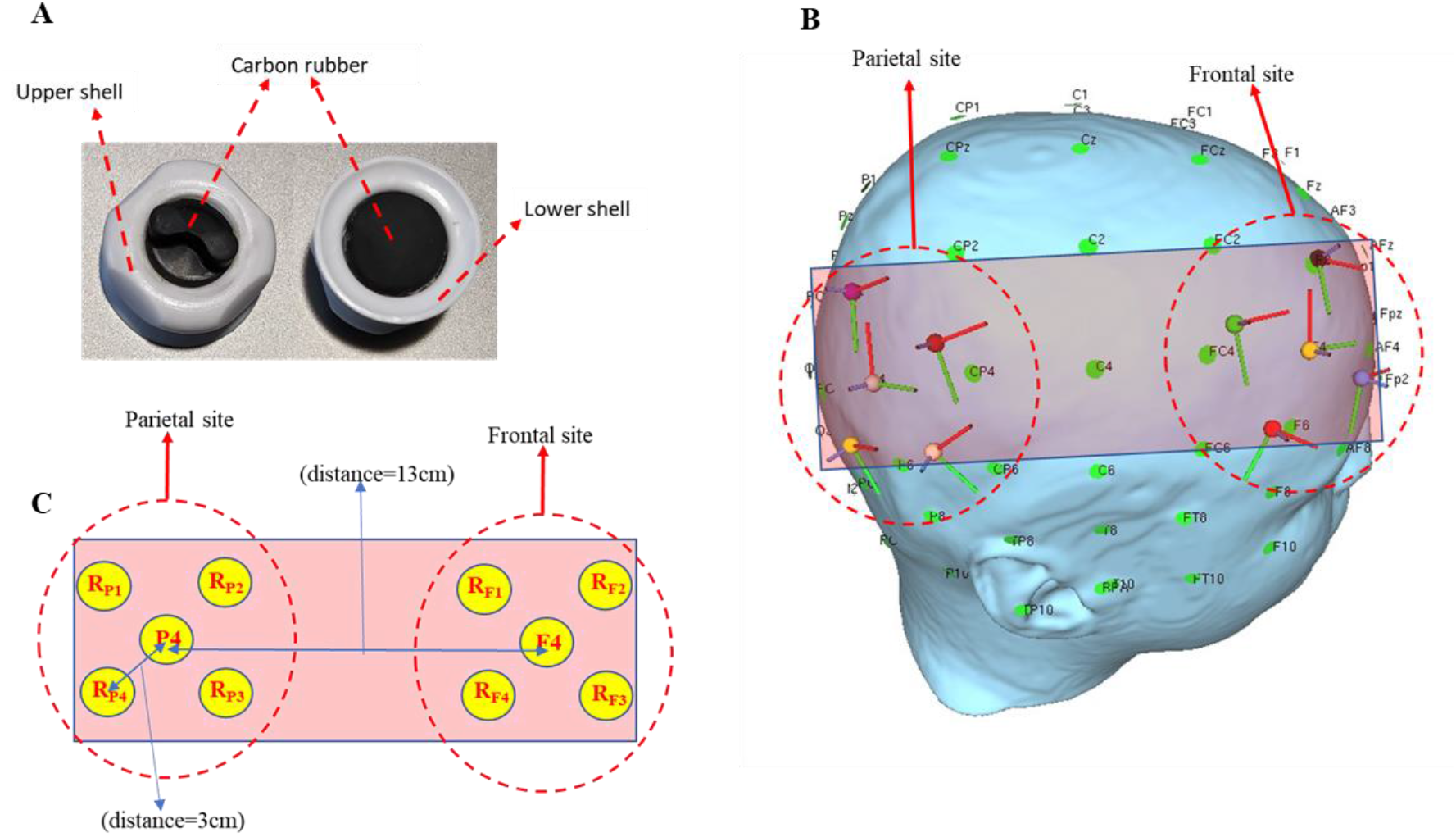
MR compatible HD electrodes and montage settings for this study. **A)** MR compatible HD electrodes; **B)** Head model of equidistance center-return (3cm) electrode placement; **C)** Plane surface of equidistance (3cm) electrode placement, with distance 13 cm between sites. To simplify analyses, we ignored the head curvature, but drew the surface of the head on which the electrodes are positioned as a plane surface, as shown in Figure 5B-C above, and the electrodes side view as shown in Figure 6 (with where the peripheral electrodes aligned in the direction of view and occluding each other being and combined into one electrode.

### 2.3 Electric field of montage

Electric field derivation in-phase and anti-phase conditions from our montage can be found in supplementary materials. Derivations show that the electric field on the in-phase condition from our montage will appear under frontal and parietal electrodes but will not appear in between under frontal and parietal electrodes. Any appearance of the electric field in between sites is the electric shunt effect (Saturnino et al., 2017). It will increase the stimulated area or decrease focality. This is not desirable if we need to focus stimulation over a specific region. Therefore, the in-phase condition is relatively safe from shunt condition. Our montage with 13cm distance between each site does not show the electric shunt effect between their sites (Figures 6A, B). Meanwhile, on the anti-phase condition, there is possibility appearing the electric shunt effect in between each site (equation 6 in supplementary). Therefore, based on equation 6, and to avoid the shunt effect, we need to pay attention to: (i) ensure sufficient distance between the return electrodes of the two sites and (ii) position the return electrodes as close as possible to their center electrode (*d*_1_ ≈ *d*_2_ ≈ *d*_3_) but do not too close to prevent too much shunting effect via the skin between center and surround electrodes (Neri et al., 2020). In our montage it is feasible to establish a return-to-center distance of 3cm, resulting in a gap of 1cm between the edges of the center and return electrodes (each of them having a diameter of 2 cm). With this montage, the return-to-center electrode distance is relatively small and the return-to-return electrode distance between frontal and parietal sites is relatively large so that the shunt effect between frontal and parietal sites in the anti-phase condition is minimized.

**Figure 6.**
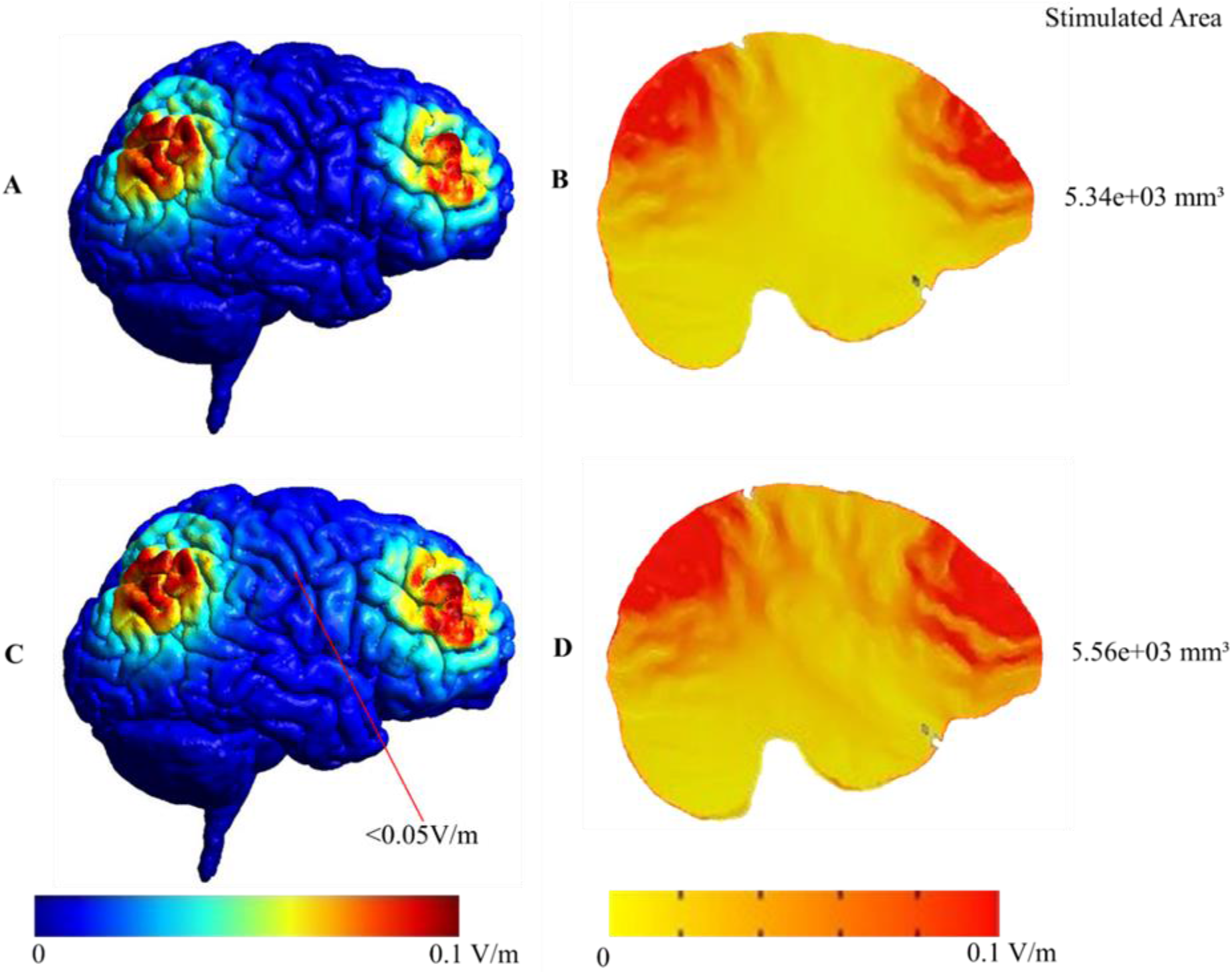
Surface- and volume-based comparison of in- and anti-phase condition. The cortical surface (surface-based electric field (EF) distribution) is calculated by SimNIBS (**A** and **C**) and could be interpolated into a NIfTI volume and applied to transformed to MNI space (volume-based electric field distribution). Therefore, we can analyze the electric field in each voxel using AFNI software (**B** and **D**). **A)** and **B)** Surface and volume-based simulation result of the in-phase condition; **C)** and **D)** Surface and volume-based simulation result of the anti-phase condition. The stimulated area in the anti-phase is 4.12 % wider than in the in-phase condition due to the electric shunt effect. In anti-phase condition, the electric shunt effect can be seen as a stronger electric field (red color) in between sites (**D**) and more electric field dots of less than 0.05V/m on the cortical surface in between sites **(C).**

From our montage, by using equation 6 and data; 2d_2_ = distance from F4 to P4 around 13cm (https://www.biosemi.com/headcap.htm), 2d_2_ = d_1_+ d_3_ = 13cm and gray matter conductivity = 0.275 S/m (Wagner, Zahn, Grodzinsky, & Pascual-Leone, 2004), we calculate the maximum electric field in the gray matter in between two sites to be 0.04 V/m. The electric field in the cortical target regions of interest in frontal and parietal cortex is provided by SimNIBS. The top percentiles of the electric field intensity in 99.9% = 9.22e-02 V/m or near to 0.1 V/m which appears on the cortical surface under the center electrode for each site (frontal and parietal) (Figure 6). It shows that the electric field obtained on cortical under each site is relatively high or the shunting effect between center and their surrounding return electrodes could be neglected. To test these hypotheses in more detail *in silico*, we simulated the electric field in the brain using SimNIBS 3.2 software (Saturnino et al., 2019; Thielscher, Antunes, & Saturnino, 2015). SimNIBS 3.2 uses the finite element mesh (FEM) method to calculate the electric field on every tetrahedron element mesh in every brain segmentation. It can be interpolated onto the cortical surface (surface-based electric field distribution) or interpolated into a NIfTI volume and transformed to MNI space (volume-based electric field distribution). Therefore, we can analyze the electric field in each voxel. SimNIBS also provides information about the focality of the stimulated area, which is defined as the grey matter volume with an electric field greater or equal to 75% of the peak value. To avoid the effect of outliers, the peak value is defined as the 99.9th percentile. The smaller the value of this volume metric, the more focal the electric field in the brain. Figures 6A and B depict the intensity of the electric field on the cortical surface, on volumetric sagittal view and the stimulated area for F4-P4 in-phase, meanwhile, Figures 6C and D show for F4-P4 anti-phase. Figures 6A, B, C, and D show the electric field is focused under frontal and parietal sites as predicted by formula 4. However, in the anti-phase condition, the electric field in between sites appears stronger in the in-phase condition, and also the stimulated area is wider than the in-phase condition (in-phase stimulated area = 5.34×10^3^ mm^3^, anti-phase stimulated area = 5.56×10^3^ mm3, percent change anti-phase to in-phase = 4.12%). This is caused by the electric shunt effect. As predicted by formula 6, the maximum shunt effect on the cortical surface is less than 0.05V/m (Figure 6C). However, the electric shunt effect for the anti-phase condition is not overly large (the stimulated area only increases by 4.12% compared to the in-phase condition), so the shunt effect can be neglected.

### 2.4 tACS-fMRI capping

Before applying the gel, we check there is no tattoo, scars, active skin irritation around electrode location. Afterward, we clean the scalp area in the electrode shell with isopropyl alcohol (IPA) using a cotton swab. This is to clean the scalp area on where the electrodes will be installed, so that dust and oil in the area will be removed to make a low impedance contact between the electrode and the scalp. We dip the cotton swab into the IPA, then swab the scalp under the electrode shell with the cotton swab evenly and gently (Figure 7A). We repeat this three or four times. Then we apply Abralyt HiCl gel to the scalp area inside the electrode’s shell, so that the amount of gel avoids a short circuiting the electrodes. If a short circuit occurs between electrodes, tACS will not work as intended. The gel must be spread evenly across the scalp inside the shell, and the level of gel should not exceed the thickness shown in Figure 7B, which is about 1 mm or the amount of 0.5 ml.

**Figure 7.**
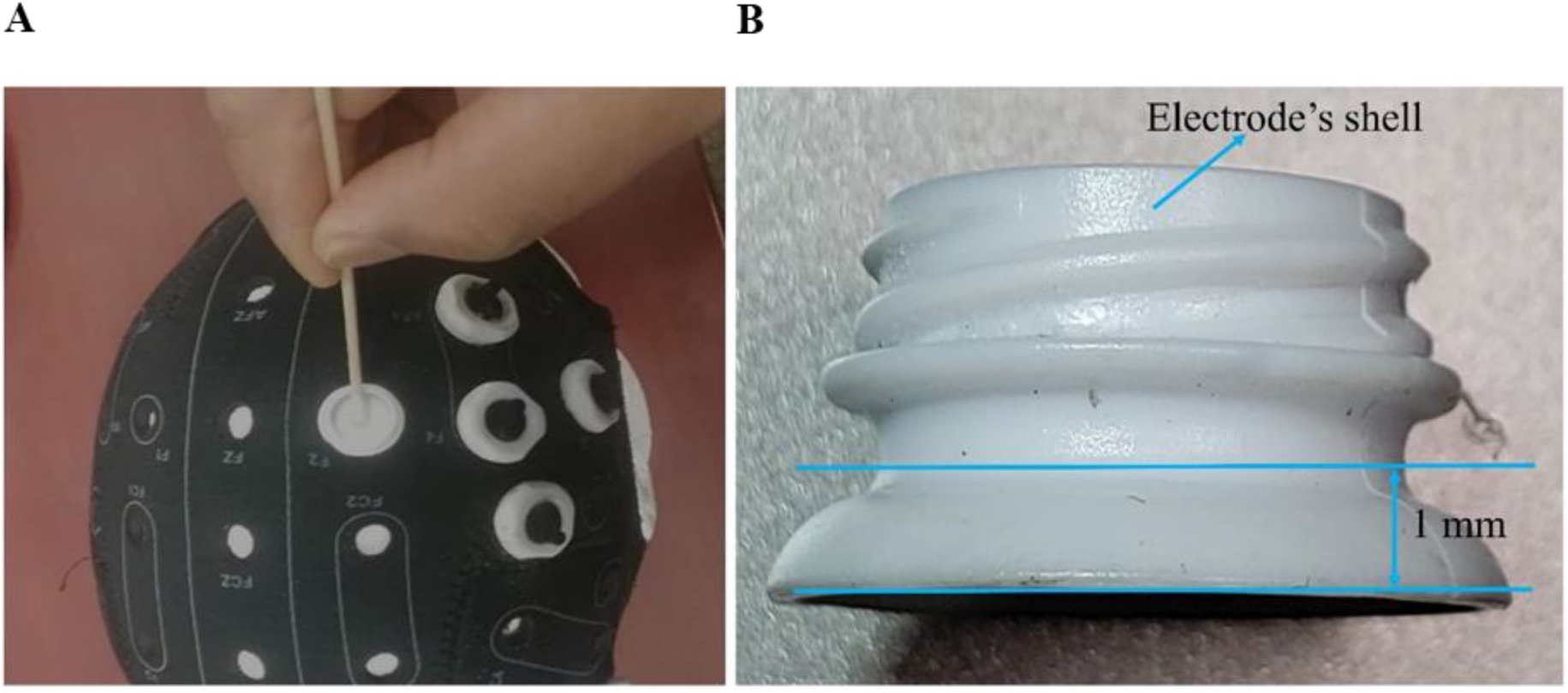
Peripheral cautions for tACS capping. **A)** Swab evenly and gently using a cotton swab with isopropyl alcohol on the scalp area inside the electrode shell. Repeat three or four times. **B)** After 10 HD shells are attached into the holes on the cap referring to the montage location, the gel is spread evenly across the scalp inside the shell. The layer of gel should not exceed 1 mm thickness or 0.5 ml gel volume to avoid excessive leakage of gel, which could make a short circuit to nearby electrodes.

### 2.5 Parameter optimization

Information processing in the brain is associated with neuronal oscillations. Furthermore, the transfer of information across various brain areas is characterized by inter-area oscillatory synchronization (Buzsáki & Draguhn, 2004; Canolty & Knight, 2010). Oscillatory synchronization of two brain regions will also temporarily increase the BOLD fMRI connectivity measures between those areas. Therefore, the fMRI connectivity between the frontal (area under the F4 electrode) and parietal sites (areas under the P4 electrode) becomes a meaningful quantitative value in measuring the synchronization level of internal brain activity. Other studies proposed that external oscillatory stimulation of cortical regions using tACS can increase internal oscillatory synchronization across brain regions and respective increases in functional connectivity measures (Cabral-Calderin et al., 2016; Kuo & Nitsche, 2012; Moisa et al., 2016; Violante et al., 2017; Weinrich et al., 2017b; Williams et al., 2017; Zoefel et al., 2018). Therefore, in this study, we try to find the optimal tACS oscillation by observing frontoparietal functional connectivity using concurrent BOLD-fMRI. Furthermore, the flow of information between brain areas may also be flexibly reconfigured through phase synchronization (Akam & Kullmann, 2014; Womelsdorf et al., 2007), and functional connectivity across distant brain regions is modulated in a phase-dependent manner (Violante et al., 2017).

Therefore, phase optimization aims to find an optimal phase difference between frontal and parietal sites, which is most improves frontoparietal functional connectivity. We use Simplex optimizer or the Nelder-Mead technique, which is a simple optimization algorithm seeking the vector of parameters corresponding to the maximum or minimum of any n-dimensional function F (x1, x2,..,xn), searching through the parameter space (Mathews & Fink, 2004; Nelder & Mead, 1965; Singer & Nelder, 2009). As our parameter space is a two-dimensional space (frequency and phase), the Simplex shape is a triangle. The main reason for choosing Simplex optimization is its ease of implementation in the real-time setting and its robustness (Huang, 2018; Price, Coope, & Byatt, 2002). The study protocol introduced in this paper is a real-time experiment (Figure 2A) with only 30 opportunities for functional connectivity calculation (2×15 stimulation blocks in training 1 and training 2). Therefore, we need an optimizer that will work sufficiently fast in real-time to find the optimum parameters. Those goals can be achieved relatively quickly by the Simplex optimizer. The Simplex optimizer requires only a triangle of tACS stimulus parameters (frequency and phase) and the fMRI functional connectivity responses estimated from a sliding window connectivity calculation. The dynamic functional connectivity between the frontal and parietal time series is calculated using the Pearson’s correlation coefficient (r) and a sliding window with 10TRs window length (1TR = 2seconds), then we normalize the distribution using Fisher’s z-transformation. The “z” values are measured for the time series within the sliding window in each step to generate the frontoparietal connectivity values over time. Then it is fed into the optimizer, which calculates the best values simultaneously for both stimulation parameters by following Simplex rules and update the tACS device for the next stimulation block. This process runs online and in a closed-loop approach which means that the optimizer uses parameter values obtained from the current stimulation block as input to update parameter values for the next stimulation block.

The 2-dimensional Simplex begins with 3 observations of the response obtained with 3 different (frequency and frontal-parietal phase difference) initial parameter settings. Therefore, the Simplex shape is a triangle. The detail of Simplex optimizer rules can be found in Nelder & Mead, 1965. The optimal parameters can be determined quickly if the initial tACS parameters are close to the optimal parameters. Therefore, prior information about the optimal parameters is important to set the initial Simplex triangle close to those parameters. Based on the result from Violante et al., 2017, our prior parameters will be around theta band (4 - 8Hz) and phase difference = 0°, as our study has similar stimulation targets (F4 and P4) and a similar goal, i.e., to improve cognitive functions measured by the subject’s performance in a 2-back task. Therefore, it makes sense to use the center of the theta band (6 Hz) and phase difference = 0° as the center of the equilateral triangle (initial Simplex parameter) with the edges (6 Hz, 5°), (10 Hz, −3°), and (2 Hz, −3°). All values for StarStim tACS input must be integer, which is why we need to round the fractional tACS parameter values to integer values. The distance from center (6Hz, 0°) of the equilateral triangle to each edge is chosen equal to be 5 (a norm of Hz-axis and degree-axis in frequency and phase coordinates) in order to give a degree of freedom for the closed-loop system to find the optimal parameters.

Validation is an important component of the optimization procedure. The intent is for measured connectivity to increase across each training run as more optimal parameters are selected. We will evaluate this at two different levels, which relate to different standards of effectiveness. In the first case, we will use a linear mixed effect model (LME). The LME model includes fixed effects of block, group, block by group interaction, and the random effect of the subject on intercept. Blocks will be modeled as a continuous variable, and a positive coefficient would indicate increasing connectivity across time, showing evidence that the optimization procedure has some effect at the group level. For clinical utility, however, the optimizer must be effective at the individual subject level. We will test this by fitting individual linear models to each run of each participant’s data. This will allow us to assess the fraction of subjects and runs in which the optimization procedure had a measurable effect.

### 2.6 Cognitive function of interest

Working memory as a key cognitive function plays a significant role in executive functions and decision making and could be impaired in different mental health disorders including substance use disorders and schizophrenia (Bickel, Yi, Landes, Hill, & Baxter, 2011; Brooks et al., 2017; Ieong & Yuan, 2017). Therefore, working memory training has been used in different treatment programs to improve clinical outcomes in different psychiatric and neurologic populations (Brooks et al., 2017; Klingberg et al., 2005; Klingberg, Forssberg, & Westerberg, 2002). Working memory has been divided into two main processes (Sauseng, Klimesch, Schabus, & Doppelmayr, 2005): (1) executive control, which manages manipulation and retrieval of information from working memory, and (2) active maintenance, which maintains the available information. The executive control function is handled by a wide region in the prefrontal cortex such as dorsolateral prefrontal cortex (DLPFC), as well as posterior and inferior regions of the prefrontal cortex. Meanwhile, active maintenance is handled mainly within the parietal cortex (Cohen et al., 1997; Prabhakaran, Narayanan, Zhao, & Gabriel, 2000). Information exchange between these two prefrontal and parietal regions which form ECN can be maintained with frontoparietal synchronized oscillatory activity in the theta range (4–8 Hz) frequency (Buzsáki, 1996; Rutishauser, Ross, Mamelak, & Schuman, 2010; Sarnthein, Petsche, Rappelsberger, Shaw, & Von Stein, 1998).

There is a growing body of evidence that appropriate external oscillatory stimulation with tACS on cortical regions may modulate internal oscillatory synchronization and functional brain connectivity (Cabral-Calderin et al., 2016; Kuo & Nitsche, 2012; Moisa et al., 2016; Reinhart & Nguyen, 2019; Violante et al., 2017; Williams et al., 2017; Zoefel et al., 2018). Therefore, we have selected working memory and one of its well-known experimental paradigms (n-back task) as the primary outcome of our online FPS tACS-fMRI brain modulation intervention. Our goal is to find the appropriately individualized parameters of external oscillatory stimulation that increase functional connectivity between frontal (F4) and parietal (P4) regions which is hypothesized to enhance working memory performance. The right frontoparietal hemisphere is stimulated according to the Violante et al., 2017 study which reported working memory enhancement within theta band (6 Hz) FPS via tACS. This finding is supported by previous evidence (Jaušovec, Jaušovec, & Pahor, 2014). Previous studies reported that stronger activity and connectivity in working memory networks significantly correlated with increasing demand in verbal N-back conditions (Dima, Jogia, & Frangou, 2014; Fedorenko, Duncan, & Kanwisher, 2013). Based on these arguments, we will compare working memory performance between optimal vs. control stimulation groups. We hypothesize that optimal stimulation is associated with higher frontoparietal synchronization and higher working memory performance than control stimulation.

## tACS-fMRI quality check and safety

### 3.1 Background

Before applying tACS-fMRI, it is necessary to verify the impact of tACS-fMRI on subject fMRI image quality and safety. Reliable and safe setups for the application of simultaneous tACS-fMRI are well known (Chaieb et al., 2014; Frank et al., 2010; Gbadeyan, Steinhauser, Mcmahon, & Meinzer, 2016; Loo et al., 2011; Poreisz, Boros, Antal, & Paulus, 2007; Williams et al., 2017); however, there is no published evidence on the safety of simultaneous tACS-fMRI with double site HD montages. We combined FPS tACS-fMRI measurements to prove that it has no aversive impact on patient safety and image quality. We aimed to: (1) to examine whether tACS stimulation induces any artifacts or increases noise on MRI/fMRI images, and (2) to conduct a tACS safety test regarding the scalp temperature under stimulation electrode during concurrent tACS stimulation during fMRI.

### 3.2 MRI Artifacts, fMRI Noise Testing Method

We use the same tACS stimulation device (Starstim R32; Neuroelectrics Barcelona SLU; Spain) inside the MRI (3T MRI scanner (Discovery MR750; GE Healthcare Systems, Milwaukee, WI) with an 8-channel receive-only head coil). Single-shot gradient-recalled echo-planner imaging (EPI) with sensitivity encoding (SENSE) is used for the scans with the parameters of FOV = 240 × 240 mm, matrix = 96 × 96 reconstructed into 128 × 128, SENSE acceleration factor R = 2, 45 axial slices with slice thickness = 2.9 mm, TR/TE = 2/0.025seconds, flip angle = 90°. The tACS montage is the same as described in section 2.1, which used 10 HD electrodes with 4 × 1 ring montage at two sites (F4 and P4 are active electrodes). F4 and P4 electrodes are selected as the main nodes of the frontoparietal network (1mA, 6Hz, 0° phase difference; Figures 4 and 5), and eight electrodes are used as return electrodes surrounding F4 and P4 with coordinates are described in section 2.1. We use three scanning protocols for different goals. The first scan is obtained without radio frequency (RF) to evaluate the impact of tACS on the EPI k-space during no-RF excitation or only noise to draw EPI images. The second is obtained with RF excitation to evaluate the impact of tACS on voxel-wise EPI images and temporal signal to noise ratio (TSNR). And the third one is to confirm tACS-fMRI safety.

We used a watermelon phantom for the first and second scans to avoid the effect of a neural activation signal modulated by tACS stimulation. The stimulation block in the first and second scan is a 20-seconds tACS stimulation ON and following by a 10-seconds no-stimulation, then the block is repeated 15 times with a 12-seconds OFF block at the beginning. The initial 12-seconds of data are excluded from the analysis to ensure a steady-state fMRI signal. Without-RF EPI scanning data on the first scan has dimension [128 128 45] and time course = 225TRs (10 × 15 = *150TRs* from Stim ON, and 5 × 15 = 75*TRs* from Stim OFF). Data is transformed by two-dimensional fast Fourier transform (2D FFT) to obtain k-space data. Since this data is collected without-RF excitation, we can evaluate the tACS influence on the MRI system noise in the receiving signal free from the sample signal, and without having a flip angle to synchronize proton magnetization phases. Therefore, phase-encoding will be encoded randomly from 0° until 360°, asynchronous with tACS, ensuring that the noise amplitude in the phase encoding direction collapses once averaged, leaving the data frequency-encoding and z-slice selected along the time course (dimension: [128 45 225]). Next, ON and OFF stimulation data are separated, and a t-test is performed between ON/OFF conditions for each frequency encoding (kx-direction) and z-slice selection. All corresponding p-values are corrected for multiple hypothesis comparisons testing using the Benjamini and Hochberg procedure for False Discovery Rate (FDR) discovery (Benjamini & Hochberg, 1995). For the voxel-wise analysis in the second scan, we performed GLM analysis on the image time-course data using 3dDeconvolve in AFNI (https://afni.nimh.nih.gov/). The regressors included a boxcar time-series for the ON period and 3rd-order Legendre polynomials to remove the low-frequency fluctuation. For TSNR analysis, it also uses EPI data images with-RF scan. First, ON and OFF stimulation data of EPI data images are separated. Then TSNR is calculated for each ON and OFF stimulation data using 3dTstat to create voxel-wise TSNR. Afterward, the mean and standard deviation of voxel-wise TSNR inside hemisphere region of interest (ROI) with radius = 10mm under frontal and parietal sites from each stimulation ON and OFF to be calculated for t-test statistics evaluation.

For the fMRI safety evaluation in the third protocol, we performed a concurrent tACS-fMRI scan for a healthy female volunteer (age 38 years) to measure temperature change on the scalp under the electrodes due to concurrent tACS stimulation during fMRI. The temperature is obtained by placing MRI compatible temperature sensors (Biopac TSD202A and Biopac SKT100C, sensitivity = 100 micro °C, sample rate = 200points/seconds) under the electrodes (P4 and F4). We used the same montage (Figures 4 and 5), fMRI parameters, and tACS parameters as in the first and second scans. We collected a baseline temperature for 2 minutes before the scanning and stimulation. Then, 2-minutes ON and OFF blocks are repeated three times to see the tACS effect in a long time period. The temperature difference between the ON and OFF periods is tested with z statistics. This study was conducted in accordance with the Declaration of Helsinki and all methods were carried out in accordance with relevant guidelines and regulations.

### 3.3 MRI/fMRI Safety Testing Results

#### 3.3.1 MRI artifacts, fMRI noise

The FDR map of the k-space analysis from the first fMRI scanning run is shown in Figure 8A. The smallest value in the FDR map from frequency encoding and z-slice selection is 0.35 (at z=19, frequency-encoding index=124), which is >0.05, meaning that tACS did not produce any significant artifact in the fMRI data. Also, on the voxel-wise analysis (with-RF), statistics on the watermelon data revealed the smallest FDR value to be 0.312, i.e., >0.05. Further analysis from the second scan, TSNR analysis looking at the ROIs under the electrodes of frontal and parietal sites shows no significant difference between tACS ON and OFF (ON: mean=10.87, SD=9.42; OFF: mean=10.73, SD=9.16, z=0.67, p=0.49), suggesting that tACS did not change TSNR. The noise caused by tACS did thus not seem to influence image quality. Also, visual inspection of voxel-wise TSNR from tACS ON/OFF indicated no tACS-related artifacts in the EPI data (Figure 8B.).

**Figure 8.**
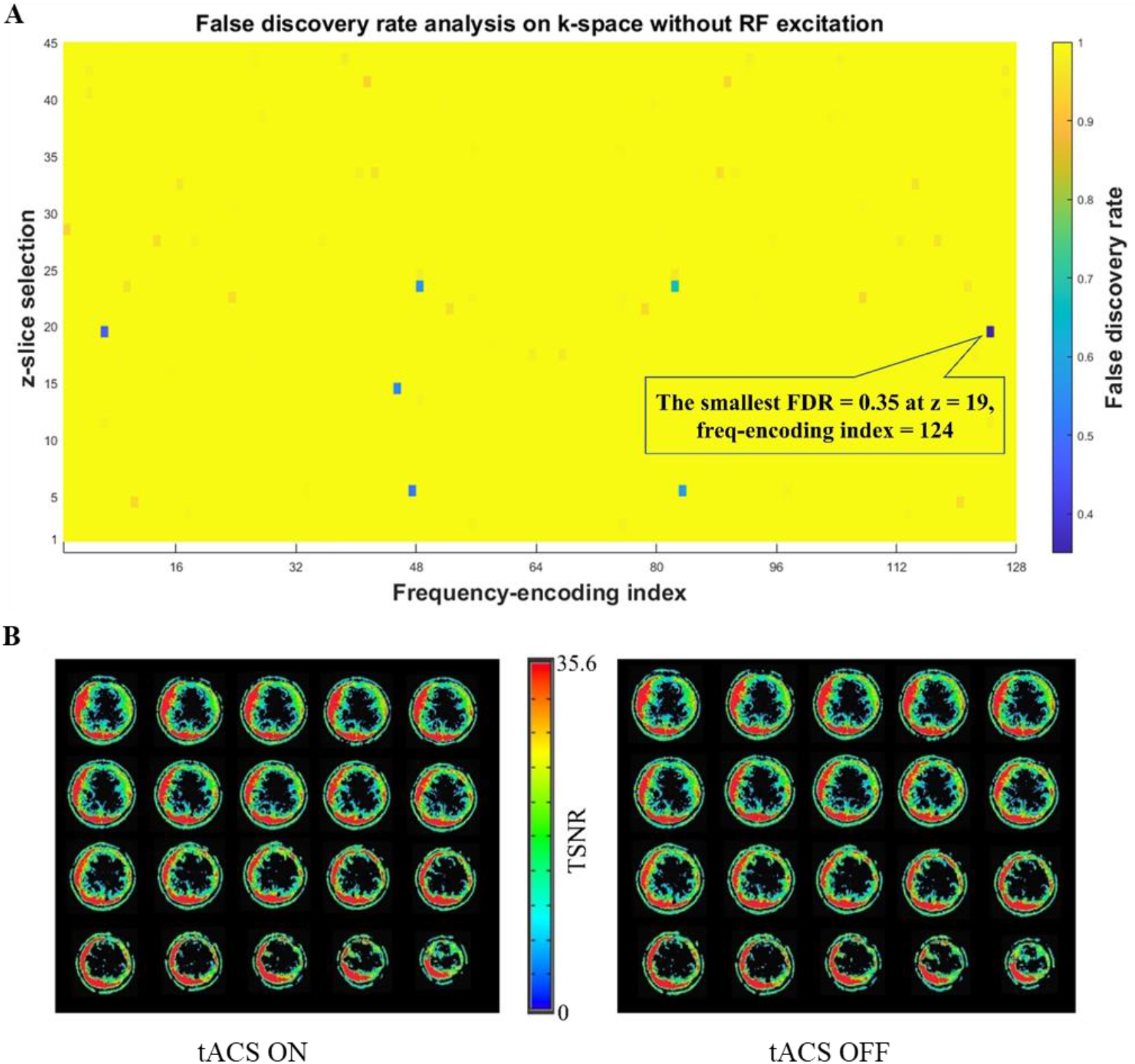
Results of the noise test influencing EPIs. **A)** FDR result/map from k-space without-RF excitation analysis. The smallest FDR value = 0.35 which is bigger than 0.05, it means tACS stimulation did not create significant artifacts. **B)** Voxel-wise TSNR from stimulation ON and OFF. TSNR analysis shows no significant difference between ON and OFF stimulation and visual inspection corroborates there is no tACS-related artifacts are observed in the EPI images.

#### 3.3.2 Temperature measurements results

The normal human body temperature range is typically observed in a range from 36.5 to 37.5 °C (Hutchison et al., 2008; Mackowiak, Wasserman, & Levine, 1992). The baseline scalp temperatures prior to scanning and tACS stimulation are stable below 33°C (F4: mean = 30.30, SD = 0.003; P4: mean = 32.22, SD = 0.05) (Figure 9A). The EPI scan did not cause a significant heating effect at the tACS electrodes (F4: mean = 30.37, SD = 0.08; P4: mean = 32.05, SD = 0.05) (Figure 9B). Moreover, the temperatures did not significantly change regardless of the tACS stimulations ON or OFF ([F4 ON: mean = 30.39, SD = 0.11; F4 OFF: mean = 30.35, SD = 0.05; z = 0.68, *p* = 0.49], [P4 ON: mean = 32.04, SD = 0.04; P4 OFF: mean = 32.07, SD = 0.05; z = −0.59, *p* = 0.56]). Furthermore, the scalp temperatures under the electrodes are below 37.5° Celsius during a 12-min EPI scan, confirming that there is no issue with patient safety in term of temperature change during tACS-fMRI in the current experimental set-up.

**Figure 9.**
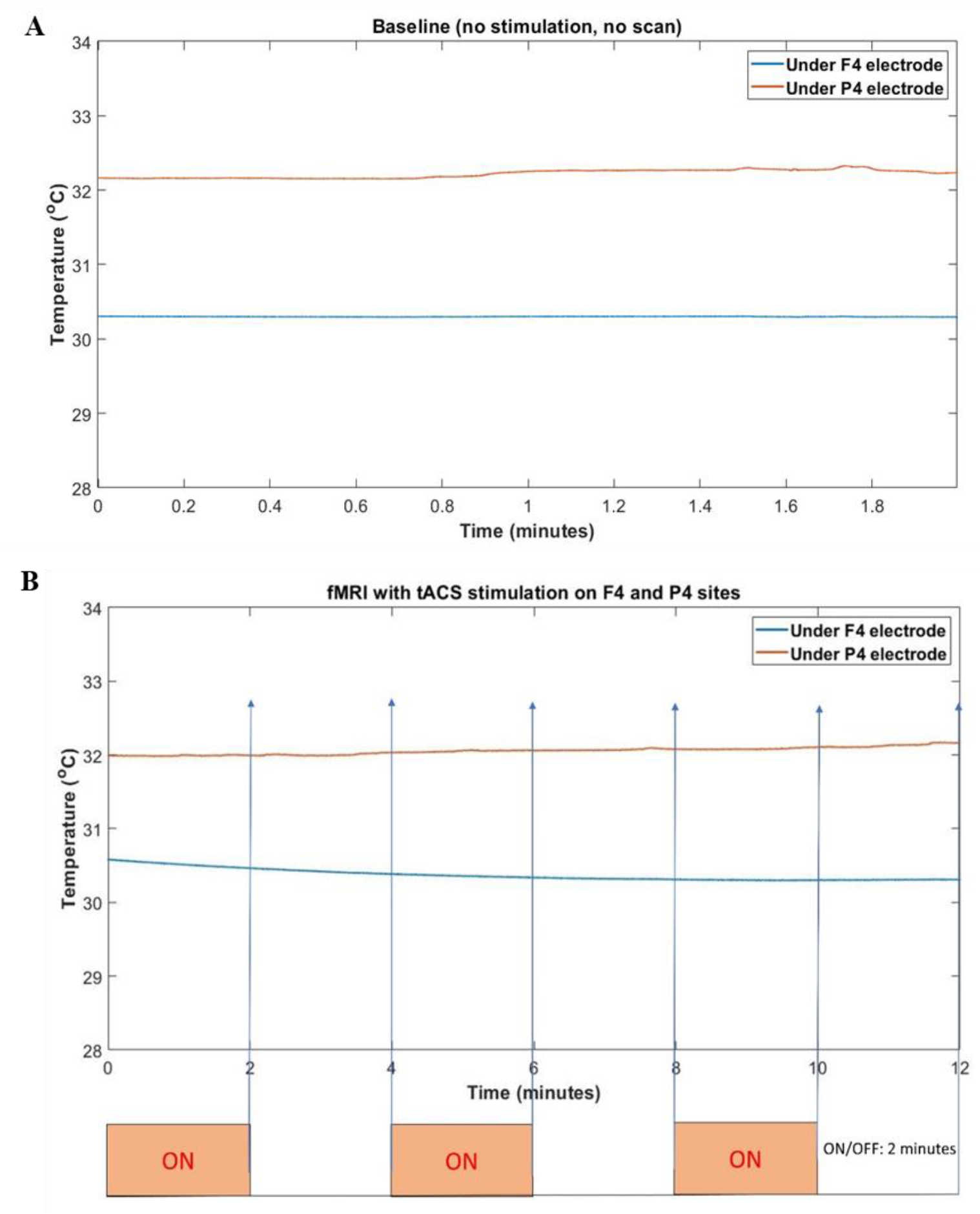
Results of the safety test influencing temperatures. **A)** Baseline temperature on the scalp at F4-P4 electrodes when there is no tACS and no scan. The baseline scalp temperatures prior to scanning and tACS stimulation are stable below 33°C; **B)** Temperature with/no tACS under F4-P4, 2 minutes ON/2 minutes OFF for 12 minutes. The temperatures did not significantly change regardless of the tACS stimulations ON or OFF. Furthermore, the scalp temperatures under the electrodes are below the upper limit human body temperature (37.5° C) during a 12 min EPI scan. It means there is no issue with patient safety in terms of temperature change during tACS-fMRI.

## Hypotheses and expected results

We aim to investigate (i) whether the closed-loop online tACS-fMRI optimization approach can optimize the tACS parameters in terms of enhancing the target functional connectivity during the training runs (the optimization phase), and (ii) whether the optimized (i.e., personalized) tACS can influence (i.e., increase) the target functional connectivity during the testing run (the testing phase), compared to a control condition. Regarding the first study aim, we hypothesize that the optimized parameter settings will be achieved along with increased target functional connectivity during the course of the training runs. Regarding the second study aim, we hypothesize that the optimized (i.e., personalized) tACS parameter settings will increase the fMRI connectivity between the tACS targets during the testing run compared to the control parameters. The first study aim cannot be tested without a control condition, since we cannot exclude non-specific changes in functional connectivity (e.g., due to boredom, habituation with MRI environment, alertness, etc.). As such a control condition is necessary even during the training run. We propose and summarized possible control conditions for this study in Figure 10.

**Figure 10.**
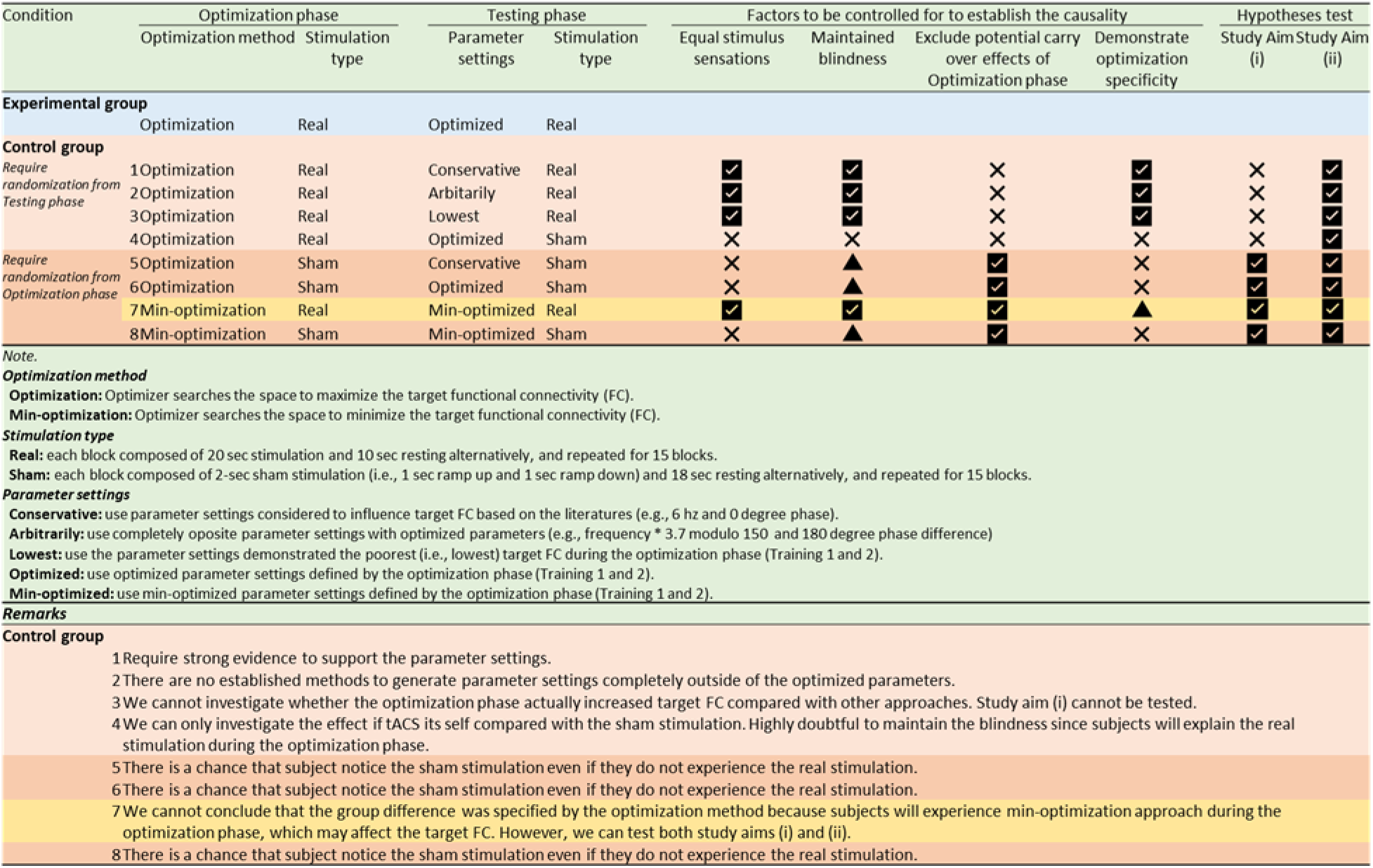
Selection of the control condition and pros and cons.

To test study aims (i) and (ii), we decided to apply the control condition described in Figure 10, condition no.7. During the optimization phase (training 1 and 2), Participants will be randomly assigned into two groups: receiving either optimized parameters (experimental group) or control parameters (control group). For the control group, the optimizer will search the tACS parameter that give the worst or lowest target frontoparietal connectivity during the training runs. We hypothesize the result of the training 1 and 2 to look comparable to the simulated data in Figure 11B. During the testing phase (testing run) participants in the experimental group will receive tACS with the optimized parameters defined by the optimization phase, while participants in the control group will receive tACS with the lowest frontoparietal connectivity parameters defined by the optimization phase. Participants will be blind to the group assignment.

**Figure 11.**
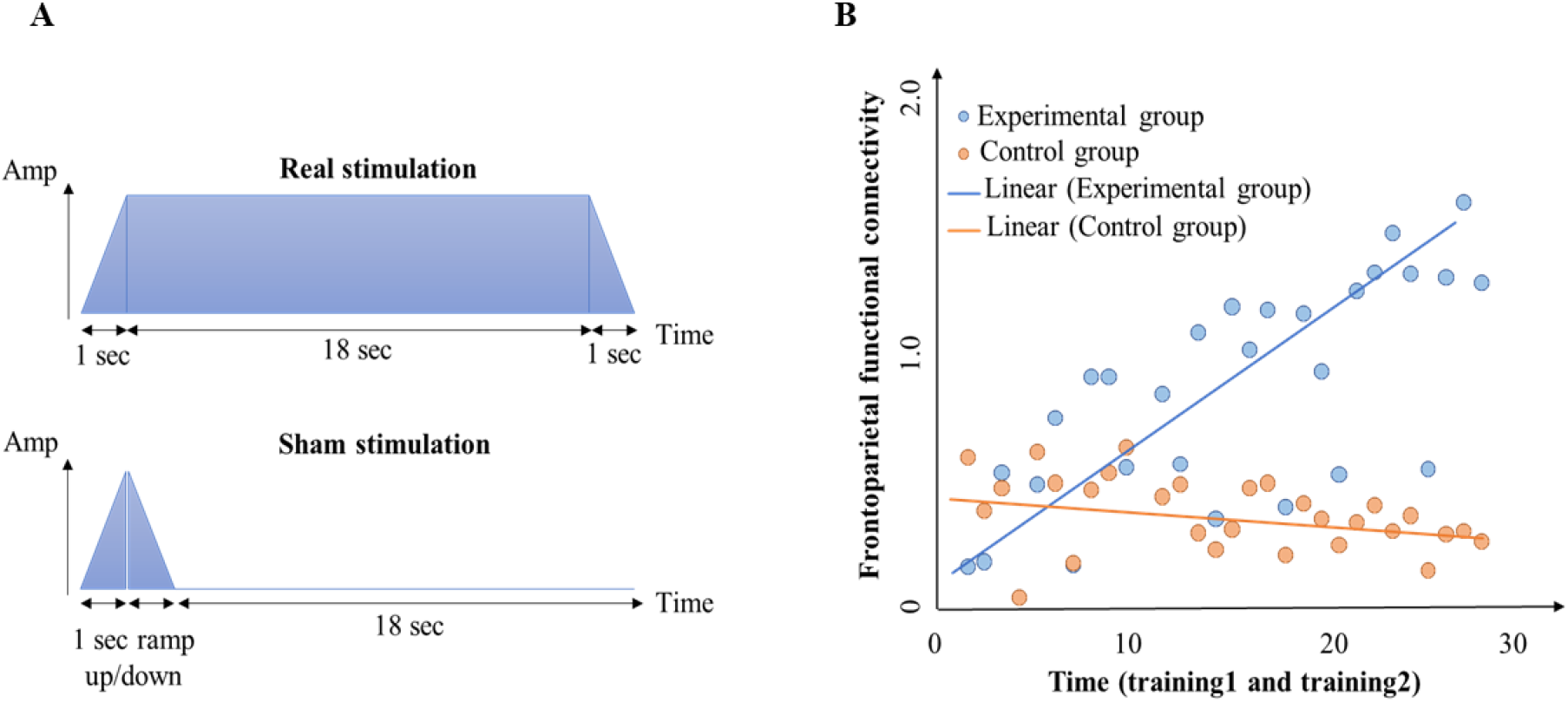
Selection of the control condition and pros and cons (cont.). **A)** graphic explanation of real and sham stimulation; **B)** hypothesized results of the optimization method between groups during the optimization phase.

Our primary outcome is the change in target functional connectivity (under the electrodes of F4-P4). The secondary outcome is the task performance during, before and after the FPS. Overall procedures of this study are described in Figure 1A. Before and after the FPS, subjects will answer mood measurements, such as Profile of Mood States Questionnaire (POMS:(McNair, Lorr, & Droppleman, 1971)) and The State-Trait Anxiety Inventory-State version (STAI-State: (Spielberger, C.D., Gorsuch, R.L., Lushene, R., Vagg, P.R., Jacobs, 1983)). During each FPS runs, we will examine whether the tACS-fMRI duration for this optimization method during the study is feasible for the participants with regard to alertness measured by Karolinska sleepiness scale (KSS:(Åkerstedt & Gillberg, 1990)), and comfort measured by visual analogue scale (VAS) (the study original). After all procedures, subjects will report potential side effects of the FPS, and also report their perception of whether they are assigned to the experimental group or the control group. Our study hypotheses are followings:

**Hypothesis 1:** The experimental group will show an increased task-related frontoparietal functional connectivity on the course of training1 and 2 compared to the control group (Figure 11B).
**Hypothesis 2:** The experimental group will show a higher task-related frontoparietal functional connectivity during the testing phase compared to the control group.
**Hypothesis 3:** The experimental group will show an accuracy improvement on the task from before the experiment (baseline) to the testing phase, compared to the control group (for which no improvement or even a decline is expected).

### 4.1 Preliminary data

The preliminary results of two subjects (experimental and control condition, respectively) are summarized in Figure 12. Figure 12A shows all blocks during training 1 and 2 for each subject. There are 30 blocks for training 1 and 2. Optimization blocks are divided into two types: successful or failed blocks. In the search for optimal parameters to get the highest or lowest frontoparietal connectivity using the Simplex optimizer, there have been several failed attempts. It is a trial-and-error algorithm, therefore, failed block to find a higher or lower connectivity than previous block cannot be counted as a path to highest or lowest connectivity and are neglected. The successful blocks track to find the highest connectivity in the experimental subject are magenta-circle-signs in Figure 12A. Meanwhile, the successful blocks track to find the lowest connectivity in the control subject are green-circle-signs in Figure 12A. The successful blocks track to find the highest or lowest connectivity are combined in Figure 12B. During the optimization phase, frontoparietal connectivity at the successful blocks in the experimental subject is consistently higher than in the control subject. It suggests that the Simplex optimizer at the end of training runs could potentially find the best tACS parameters in order to make the highest (for the experimental subject) and lowest (for the control subject) frontoparietal connectivity. The testing run also confirmed this result as shown on Figure 12C. From 15 blocks in the testing run, the normalized frontoparietal connectivity of the experimental subject is higher compared to the control subject [experimental subject: mean = 0.70, SD = 0.29; control subject: mean = 0.42, SD = 0.33; t(28) = 2.38, *p* = 0.025]. Within-subject analysis of the working memory test (2-back task) is also performed. We compare the percentage of 2-back task accuracy in testing run vs baseline (before the FPS). Figure 12D shows that the experimental subject improved in accuracy on the 2-back task during the testing run compared with the baseline, while the control subject did not show any improvement (experimental subject accuracy improvement = 8.08%; control subject accuracy improvement = 0.91). These preliminary results provide evidence for the feasibility of the protocol; however, the efficacy of the intervention should be finally tested after data collection for the entire sample of experimental and control subjects.

**Figure 12.**
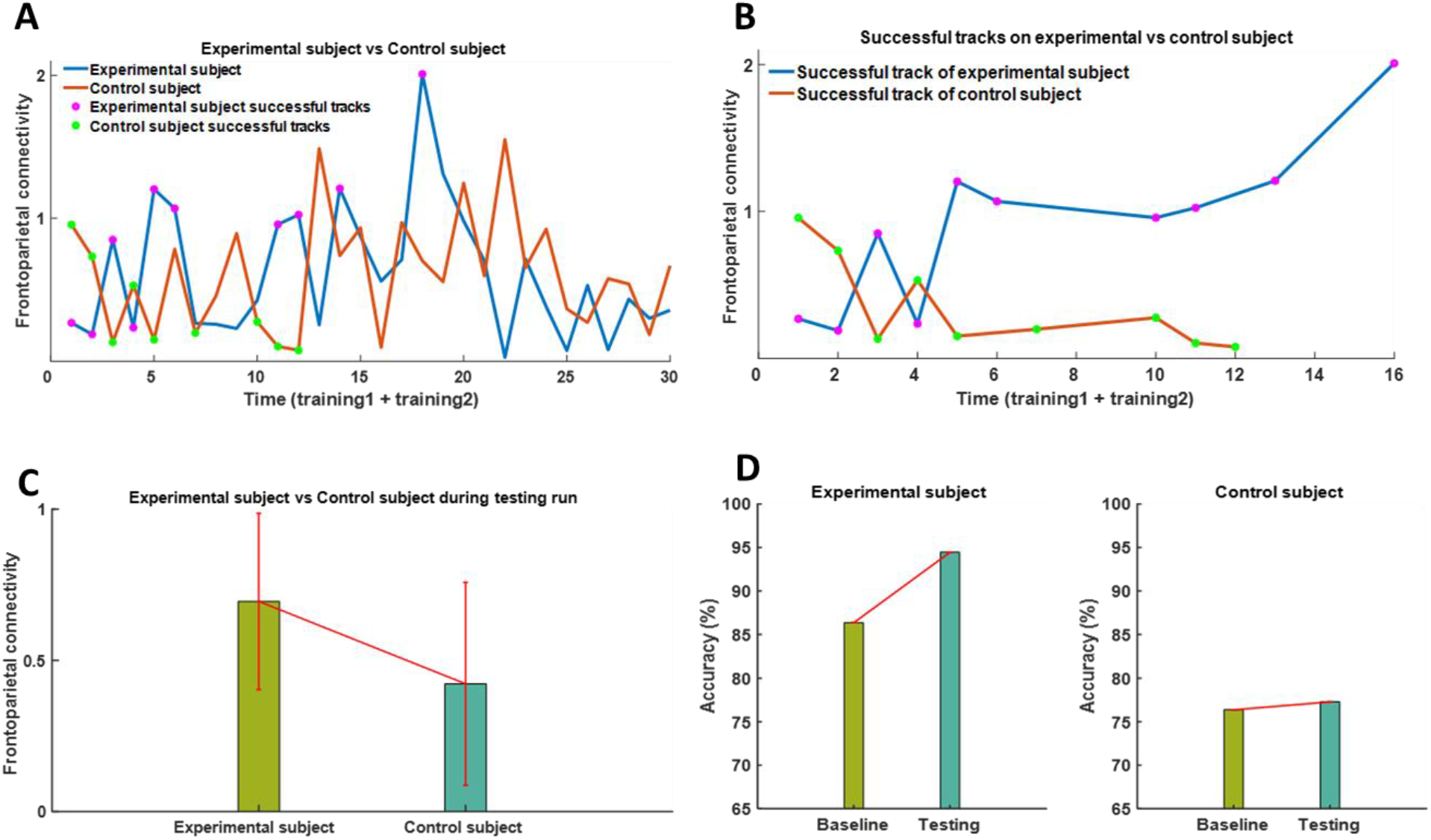
Preliminary results of the closed-loop tACS-fMRI protocol from two sample of the experimental and control subjects. **A)** Blocks on training 1 and 2 for each subject. There are 30 blocks for training 1 and 2. Optimization blocks could be divided into two types; successful or failed blocks. Failed attempts could not be counted as a path to highest or lowest connectivity, so it must be discarded. The successful blocks track to find the highest connectivity on the experimental subject are magenta-circle-signs, meanwhile the successful blocks track to find the lowest connectivity on the control subject are green-circle-signs; **B)** The combination of successful blocks track to find the highest or lowest connectivity. The experimental subject vs. the control subject has a bigger gap in frontoparietal connectivity over a time course. It means the Simplex optimizer at the end of training runs can find the best tACS parameters in order to make highest and lowest frontoparietal connectivity on the experimental subject and control subject; **C)** On testing run, The experimental subject showed significant higher the frontoparietal connectivity rather than control subject; **D)** Within-subject analysis shows that experimental subject can improve accuracy to answer correct in 2-back task test before-vs during-testing run rather than control subject [experimental subject accuracy improvement = 8.08%; control subject accuracy improvement = 0.91%].

## Potential future works

The innate temporal connectivity fluctuation associated with the fluctuation of attention during the n-back task may challenge the original Simplex optimization algorithm. The algorithm relies on the frontoparietal functional connectivity, which also might be influenced and fluctuated by individuals’ motivation to perform the 2-back task. Therefore, we need to modify the original Simplex rules if we face strongly fluctuating connectivity responses. There are potentials to elaborate the Simplex optimizer with other artificial intelligence methods. Artificial intelligence based on the previous responses could then predict the normal connectivity response for the next stimulation step that is chosen by the Simplex optimizer. From that information, we can determine whether or not the subject’s actual response in the subsequent stimulation fluctuates too much. If there is too much fluctuation, we can give an alert to the subject monitor that alarms the subject to focus and pay attention to perform the 2-back task; then the stimulation with the same parameters is performed once again to get a better functional connectivity response.

Another future potential application of this approach will be a combined tACS-fMRI neurofeedback training for substance use disorder or other psychiatric disorders. Generally, while tES is a non-invasive neuromodulation technique based on weak current stimulation over the scalp, neurofeedback is another form of non-invasive neuromodulation approach based on an individual’s learning process which helps to change brain activities by monitoring and controlling feedback signals from their brain physiological measurement (e.g., EEG or BOLD signal changes in fMRI). We can extend our current research into fMRI-neurofeedback enhanced by tACS, by presenting individuals fMRI BOLD signal changes during the 2-back task with the enhancement of individually optimized tACS. By enhancing fMRI-neurofeedback with tACS, we may be able to see additional treatment benefits for those who are not able to respond to fMRI-neurofeedback.

## Conclusion

We introduced an online frontoparietal stimulation closed-loop tACS-fMRI protocol and reviewed the available evidence to establish the optimization method of tACS stimulation parameters (i.e., electric current frequency and phase difference) utilizing simultaneous tACS-fMRI. We described the potential block design for the optimization process, tACS-fMRI equipment settings including HD electrodes and montages, and the optimization algorithm. Simulation analysis shows that focality of stimulation is observed under each frontal and parietal site during different phase conditions. Furthermore, by the specific return electrode placement, we can reduce the shunt effect of different phase stimulations to minimal values (stimulated area only increases 4.12% in the anti-phase stimulation compared to the in-phase stimulation). Also, we suggested the Simplex optimizer (Nelder-Mead technique) because it is a simple optimization algorithm, and the calculations are performed rapidly, which is suitable for real-time closed-loop experimental settings. Simplex optimizer is fairly robust to find the best parameters for the maximal response if the connectivity response is quite stable in time (Barton & Ivey, 1996). Moreover, we suggest that utilizing a task-based scan (e.g., 2-back task) as a solution to increase the effect of stimulation and reduce the instability of functional connectivity during the course of stimulation. We also conducted a safety test for this proposed protocol, and we reported that our tACS-fMRI setting will not cause any adverse heating effects or image artifacts. There are still many questions remaining with respect to the best methodological parameters for this novel integration of tES-fMRI brain modulation that should be tested empirically in future studies.

## Author Contributions

B.M., H.E., J.B., M.M., A.T., and R.K. designed the study. B.M., A.T., and J.S., collected the data. B.M. performed simulations and data analyses under H.E., M.M., J.B., S.C., and R.K. supervision. B.M. wrote the paper with input from H.E., T.O.B., D.S., M.M., R.K., G.S., A.T., A.R., S.C., J.B., and M.P. All authors (B.M., H.E., T.O.B., D.S., M.M., R.K., G.S., A.T., J.S., A.R., S.C., J.B., and M.P.) contributed in manuscript preparation. All authors (B.M., H.E., T.O.B., D.S., M.M., R.K., G.S., A.T., J.S., A.R., S.C., J.B., and M.P.) agreed on the final manuscript before submission.

## Acknowledgments

We are grateful to Julie Arterbury, Bill Alden, Julie DiCarlo, and Greg Hammond for helping with MRI and tES-fMRI scanning.

## Supplement

F4 and P4 electrodes in the 10-20 system are the centers of stimulation, with the current function of *F*4 = *A* × sin(2*π* × *freq*_*F*4_ × *t* + *phase*_*F*4_), and of *P*4 = *A* × sin(2*π* × *freq*_*P*4_ × *t* + *phase*_*P*4_). To reduce space complexity in finding optimal parameters and to reduce training time, the electric current ‘*A*’ will not become a parameter that will be searched by the optimizer, but it will be fixed to 1 mA-peak value. The current function of each of the F4 returning-electrodes is such that: 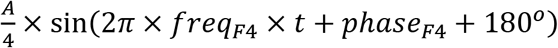. The current is divided by 4 and a phase of 180° is added in order to fulfill Kirchhoff’s law. Likewise, the electric current function on each of the P4 returning-electrodes is that: 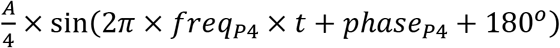. To calculate the electric field on the cortex, we use the method described by Saturnino et al., 2017. Once the electric current is applied through electrode on the scalp, then the electric field at position p inside the head and at time point t is determined by the product of the spatial component ***E***(p) and the time course of the electric current *I*(t) injected into the active channel (equation 1):

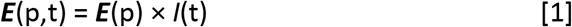

Because we have 10 electrodes which emit the current mentioned above, the total electric field at the point p as described in equation 2:

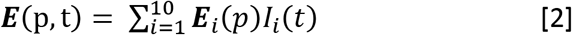

For a simpler analysis, we ignore the head curvature and draw the surface of the head in which the electrodes are positioned as a plane surface, as shown in Figures S1B and C, and the electrodes side view as shown in Figure S1A (with where the peripheral electrodes aligned in the direction of view and occluding each other being and combined into one electrode). In equation 1, the spatial component ***E***(p) is inversely proportional to conductivity (*κ*) at point p and the square of the distance (*r*) between the electrode to point p or 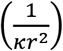. If the *phase*_*F*4_ is equal to *phase*_*P*4_, termed *in-phase* condition, then the electric field at p under P4 on the y-axis, and at time *t*_0_ using equation 2 can be written as equation 3:

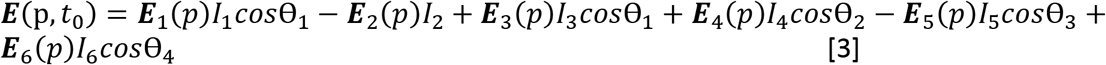

we know, ***E**_i_*(*p*) is proportional to 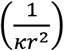, for simplification, we assume κ =1 at point p, and *I*_1_ = *I*_3_ = 0.5*I*_2_, *I*_4_ = *I*_6_ = 0.5*I*_5_, and *I*_2_ = *I*_5_ = *I_o_*. If *r*_2_ is the distance from electrode P4 to point p, then equation 3 can be written as equation 4:

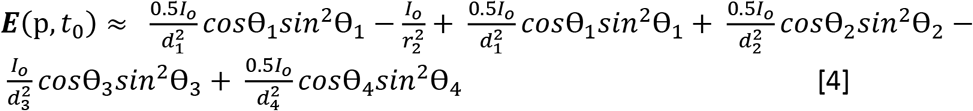

Note components: 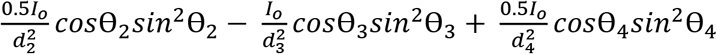 can be neglected because they are closed to zero since θ_2_ ≈ θ_3_ ≈ θ_4_ ≈ 90° or *r*_2_ is small, then *cos*θ_2_ ≈ *cos*θ_3_ ≈ *cos*θ_4_ ≈ 0.

Therefore, 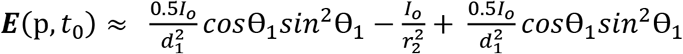 and is only influenced by electrodes above the cortex. Likewise, when we analyze along the x-axis, the electric field in in-phase condition will be dominant from electrodes above the cortex surface. If we put the p position in between frontal and parietal (Figure S1B), the electric field on the y-axis in that point is:

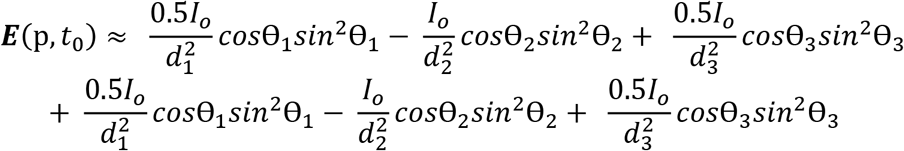

or

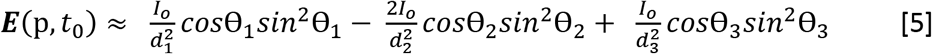

Once again if θ_1_ ≈ θ_2_ ≈ θ_3_ ≈ 90° or if *r*_2_ is small, then ***E***(p, *t*_0_) ≈ 0. The electric field along the x-axis is also 0 since all x-component are cancelled each other. Thus, in the in-phase condition, there is no electric field in any volume between frontal and parietal electrodes. Therefore, it can be concluded that the electric field on the in-phase condition from our montage will appear under frontal and parietal electrodes but will not appear in between under frontal and parietal electrodes. Then, what is the electric field in between sites if we change into anti-phase condition? It is anti-phase in the condition when *phase*_*F*4_ and *phase*_*P*4_ differ by 180°. The electric field generated from every electrode at the time *t*_0_ for anti-phase is illustrated in Figure S1C. From Figure S1C, we derive the electric field at point p and time *t*_0_ along x-axis such that:

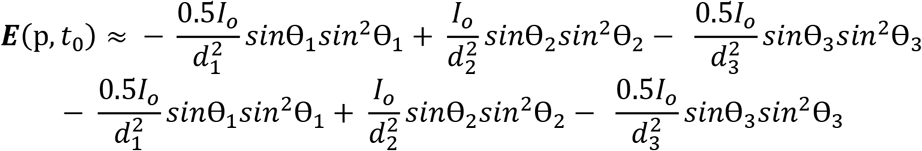

or

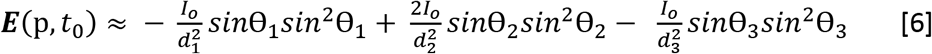

If θ_1_ ≈ θ_2_ ≈ θ_3_ ≈ 90° or *r*_2_ is small, then 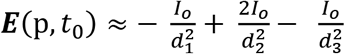, and 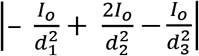 could be larger than 0 if *d*_1_ ≪ *d*_2_ < *d*_3_, *or d*_1_ ≉ *d*_2_ ≉ *d*_3_. The electric field along y-axis = 0, caused by every component is cancelled each other.

**Figure S1.**
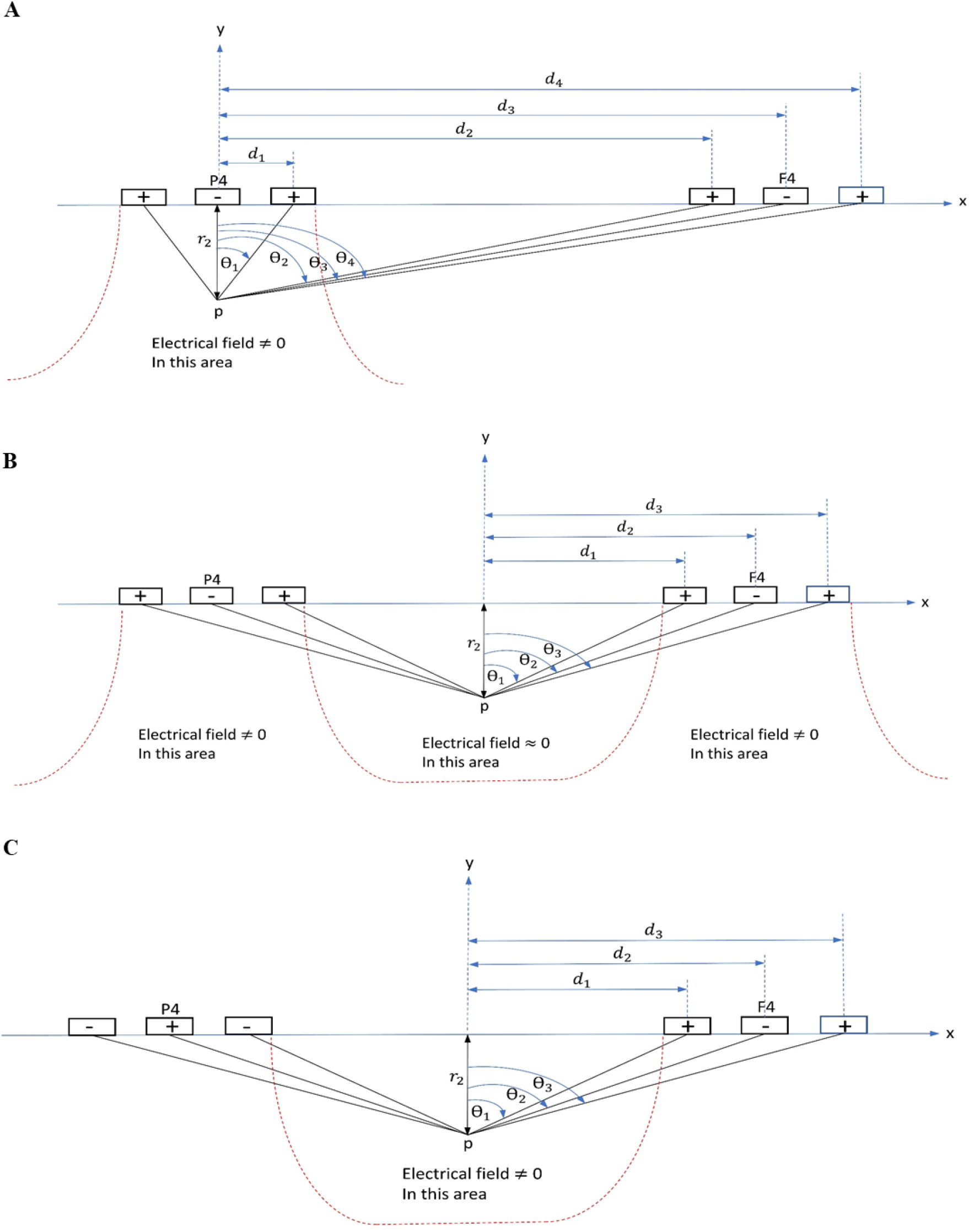
The electrodes from the sagittal side view. *d_1_, d_2_, d_3_, d_4_* and *θ_1_, θ_2_, θ_3_, θ_4_* have two points of view; [1] distance and skewed angle from the center electrode to return electrodes in frontal (F4) and parietal (P4) sites which is related to point **A**, [2] distance and skewed angle from the center in between sites to each electrode (F4, P4 and their returns) in frontal (F4) and parietal (P4) sites which is related to point **B**. **A)** In the in-phase condition, the electric field at point p is dominant from electrodes above its point p; **B)** In the in-phase condition, the electric field at point p position between frontal and parietal is ≈ 0; **C)** The electric field generated by electrodes in anti-phase condition. The electric field at point p position between frontal and parietal is ≠ 0. r_1_ and r_2_ are the distance from the scalp to point p inside the brain.

